# Identification of Differentially Expressed Genes and associated pathways common to Eyelid and Non-Ocular Basal Cell Carcinoma to understand the Molecular Biology of BCC

**DOI:** 10.1101/2021.05.01.442022

**Authors:** Pranjal Vats, Perumal Jayaraj, Seema Sen, Shefali Dahiya, Vanshika Mohindroo

## Abstract

**Background:** Eyelid BCC accounts for more than 90% of Eyelid malignant neoplasms. Various aberrant signalling pathways and genes in Non-Ocular BCC have been found whereas Eyelid bcc remains elusive.

**Objective:** This study aims to find the common DEGs of Eyelid and Non-Ocular BCC using bioinformatic analysis and text mining to gain more insights into the molecular aspects common to both BCC non-ocular and Eyelid BCC and to identify common potential prognostic markers.

**Material and method:** The Gene Expression profiles of Eyelid BCC (GSE103439) and Non-Ocular BCC (GSE53462) were obtained from the NCBI GEO database followed by identification of common DEGs. Protein-Protein interaction and Pathway Enrichment analysis of these screened genes was done using bioinformatic tools like STRING, Cytoscape and BiNGO, DAVID, KEGG respectively.

**Results:** A total of 181 genes were found common in both datasets. A PPI network was formed for the screened genes and 20 HUB genes were sorted which included CTNNB1, MAPK14, BTRC, EGFR, ADAM17. Pathway enrichment of HUB genes showed that they were dysregulated in carcinogenic and apoptotic pathways that seem to play a role in the progression of both the BCC.

**Conclusion:** The result and findings of bioinformatic analysis highlighted the molecular pathways and genes enriched in both Eyelid BCC as well as Non-Ocular BCC. The identified pathways should be studied further to recognise common molecular events that would lead to the progression of BCC. This may provide a window to explore the prognostic and therapeutic strategies common to both BCC.

## Introduction

Eyelid Basal Cell Carcinoma (BCC) is a common but unpredictable condition, accounting for over 90% of malignant eyelid neoplasms.^1^ Recurrence of Eyelid BCC is commonly seen and is considered to be more aggressive and infiltrative than the primary tumour but rarely metastasizes. The worse prognosis of the disease leads to deep invasive tumours and inadequate treatment.^2^ Its onset is majorly due to mutations caused by Ultraviolet radiation so it’s observed that lower eyelids are generally more at risk due to direct exposure to the sun.^3,4^ Prolonged exposure to UV has been the cause of TP53 mutations in more than 50% of Ocular BCCs.^5^ Ocular cancer refers to cancer of surface of eye and eyelid. It may implicate inside of eye by cancer cells growth in uvea. The ultraviolet radiation causes mutations in DNA which gets transcripted to mRNA and hampers the working of different genes by either upregulating or downregulating their expression.^4^ This DNA damage has also been reported in the “Hedgehog pathway” which is responsible for cell differentiation. Besides these aggressive histological forms of BCC have also shown aberrant expression of genes like Ki-67 Bcl-2, PTCH1, GLI1, SHH, SMO.^6,7^

BCC of the eyelid comprises more than 20% of basal cell malignancies occurring on the head and neck, both being regions of Non-Ocular BCC.^1^ Non-Ocular BCC is an amelanotic malignancy that also occurs due to the mutations caused by UV rays that produce mutagenic photoproducts in DNA leading to tumour formation.^8^ The most affected pathways in Non-Ocular BCC are the hedgehog and hippo pathway along with some secondary pathways like IGF-PI3K-AKT, EGFR-MEK-ERK and target mammalian rapamycin pathway (mTOR). Besides this; mutations in genes like MYCN, PPPC, SK19, LATS1, ERBB2, PIK23C, N-RAS, K-RAS, H-RAS, PTPN14, RB1, FBX7, S6K1 are also responsible for Non-Ocular BCC.^9– 12^

Lately, the use of computational gene network analysis along with text mining methodology has provided us with means to explore more about the occurrence of BCC eyelid. The aim was to study the networking of genes that are involved in underlying pathways and the classical pathways as well as common to both BCC of eyelid and Non-Ocular region. Therefore, the relative study of Non-Ocular BCC and BCC eyelid using a bioinformatics approach led to the identification and analysis of Differentially expressed genes and potential molecular pathways which can be used as biomarkers and further be investigated for their therapeutic practicality.

## Material and Method

### Microarray Data

Gene expression profiles of Eyelid BCC (GSE103439) and Non-Ocular BCC (GSE53462) were obtained from the National Center of Biotechnology Information (NCBI) GEO database (https://www.ncbi.nlm.nih.gov/geo/).GEO profile (GSE103439) accommodated 6 samples; including 2 control samples and 4 tumour patients who were all females and age varied from 40 to 85 years. The dataset GSE53462 comprised 21 samples; including 16 tumour patient samples and 5 control samples which were obtained from 19 females and 2 males in this study. (Fig. 1a)

**Fig 1.**
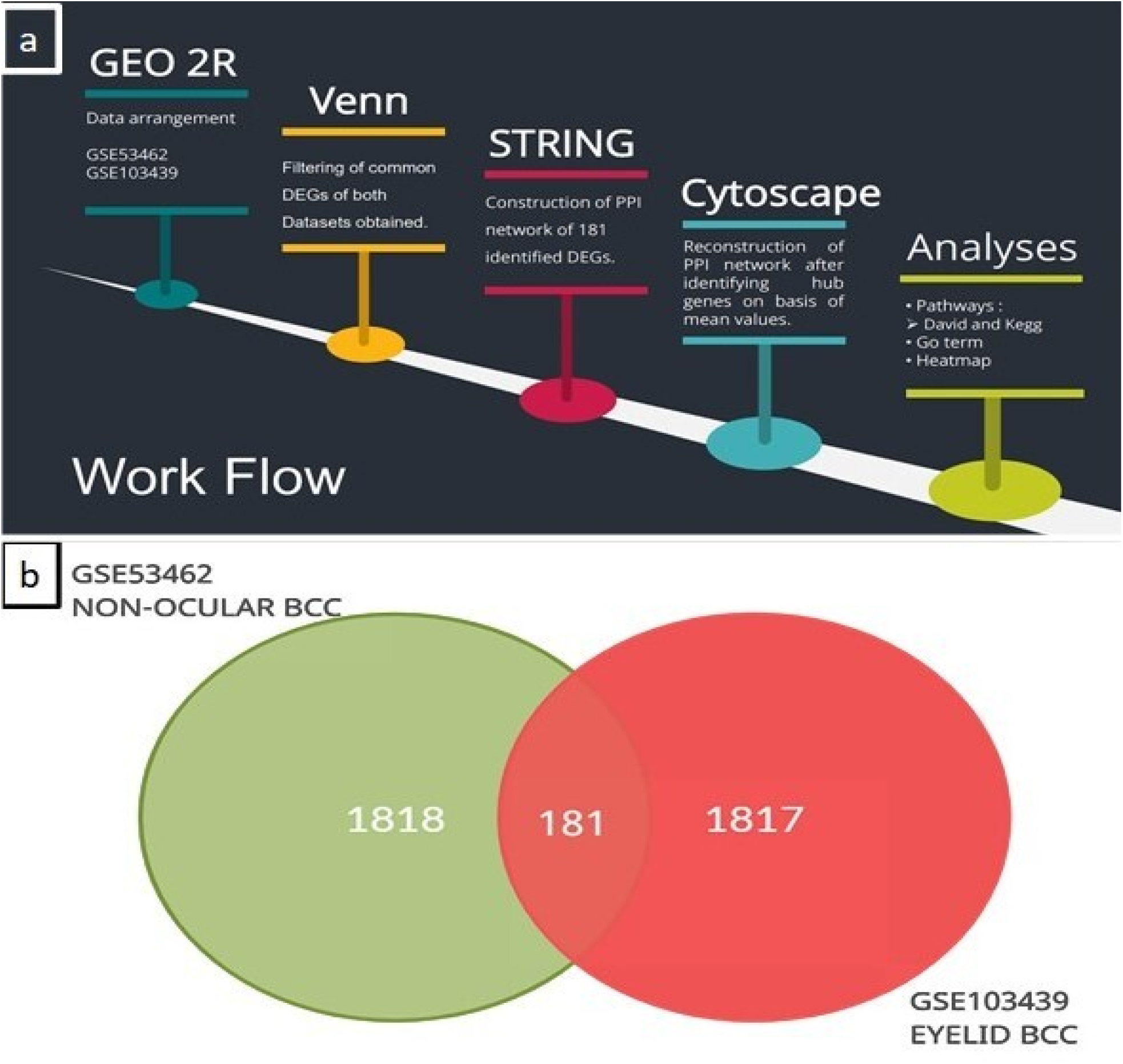
(a) Pictorial representation depicting the work flow of analysis. (b) A Venn plot of differentially expressed genes (DEGs) in two datasets GSE103439 (Eyelid BCC) and GSE53462 (Non-Ocular BCC) below adj. P-value 0.05, revealing 181 common genes in two Subtypes of BCC on the basis of location.

### Data preprocessing and identification of DEGs

Normalisation of the raw data (Fig. S1) was done using RStudio 1.3.1056 and after removing the duplicate values from the datasets, top 2000 genes were taken from both the datasets; adj. p-value of < 0.05 and common DEGs were obtained using Venn diagrammatic representation (http://bioinformatics.psb.ugent.be/webtools/Venn/). (Fig. 1b)

### Construction of PPI Network, Module analysis and identification of HUB Genes

STRING was employed on the filtered common DEGs from both datasets to construct a network based on interaction evidence and strength of data support. A combined score of >0.9 was assigned as cut off standard. This network was reconstructed in Cytoscape 3.8.0 Software to get the central network using the plug-in MCODE; cut off parameters were set as MCODE score >3 and node numbers >4 and significant hub genes were ranked using cytoHubba plug-in on grounds like degree, bottleneck, eccentricity and closeness.

### Enrichment analysis of DEGs

To further explore and look into the functions of Top 20 DEGs, pathway enrichment analysis was done using KEGG (Kyoto Encyclopedia of Genes and Genomes) (https://www.genome.jp/kegg/pathway.html), DAVID (https://david.ncifcrf.gov/home.jsp) and BiNGO from Cytoscape. The various criteria used for GO analysis were-Hypergeometric test for statistical test, Benjamini and Hochberg False Discovery Rate (FDR) correction as mode of multiple testing correction and a significance cutoff of 0.05. GO_Biological_Process was used as an ontology file to determine and annotate the significant biological pathways in which all the DEGs were either taking part directly or indirectly to activate/deactivate the pathways.

## Results

### Data processing and Identification of common DEGs

Following screening with the criteria of P<0.05 and fold-change > 2, a total of 1817 DEGs were identified. Thereafter, a total of 181 common DEGs were filtered using Venn and incorporated into this study. (Fig. 1b)

### PPI Network and HUB Genes

According PPI network from STRING online tool; 43 genes were found in close network among which CTNNB1, EGFR, GNG2, BTRC and MAPK14 had a higher degree of connectivity whereas protein that didn’t interact were removed to get the final network. PPI network of HUB genes having 43 nodes and 54 edges is presented (Fig. 2). With the help of the MCODE plugin, the top 2 sub-networks with most importance were analysed (Fig. 3a and 3b). Following calculations by cytoHubba, screening of HUB genes using values of degree, closeness, betweenness and bottleneck was done. (Fig. 3c) (Table 1)

**Table 1:-.** Topological parameters of the hub genes.

| Gene | Degree | Betweenness | Closeness | Bottleneck | Regulation |  |
| --- | --- | --- | --- | --- | --- | --- |
|  |  |  |  |  | Ocular | Non - Ocular |
| CTNNB1 | 10 | 518.6666667 | 18.81666667 | 32 | Up | Up |
| EGFR | 7 | 443 | 16.75 | 22 | Down | Up |
| BTRC | 6 | 260 | 15.03333333 | 6 | Down | Down |
| MAPK14 | 6 | 87.66666667 | 15.48333333 | 7 | Down | - |
| GNG2 | 6 | 172 | 14.78333333 | 8 | Up | Down |
| AR | 5 | 106 | 14.98333333 | 3 | Up | Down |
| CDC20 | 5 | 60 | 12.16904762 | 2 | Up | Up |
| PRKCB | 3 | 265 | 13.86666667 | 8 | Up | Down |
| ARHGEF7 | 2 | 60 | 11.51666667 | 2 | Down | Down |
| ADAM17 | 2 | 168 | 10.88333333 | 4 | Down | Up |

**Fig 2.**
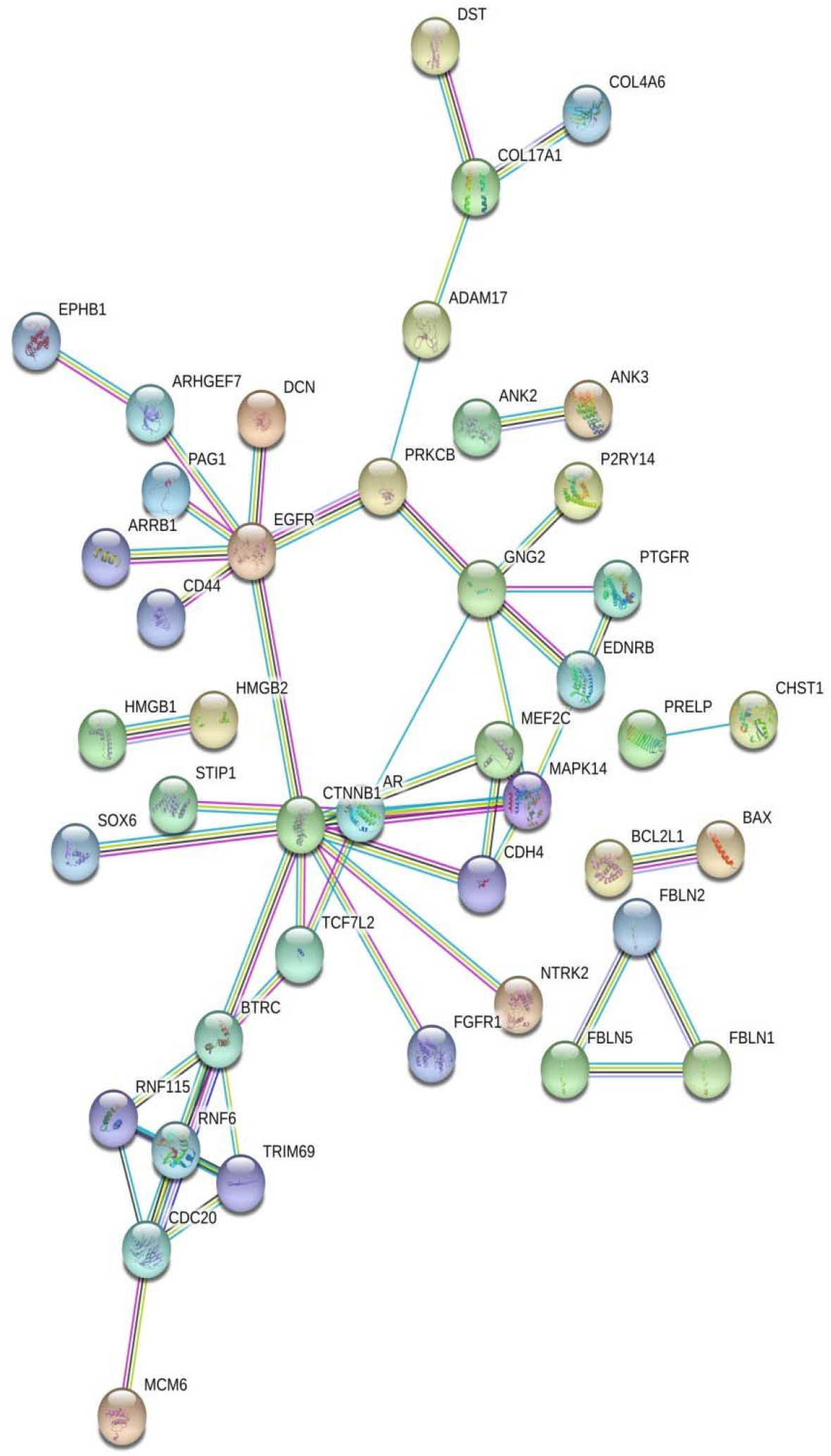
PPI network of 43 common Differentially expressed genes in STRING for evidence analysis. Each node in the given map represents a protein and the interactions made by each protein depends on highest confidence value (0.900)

**Fig 3.**
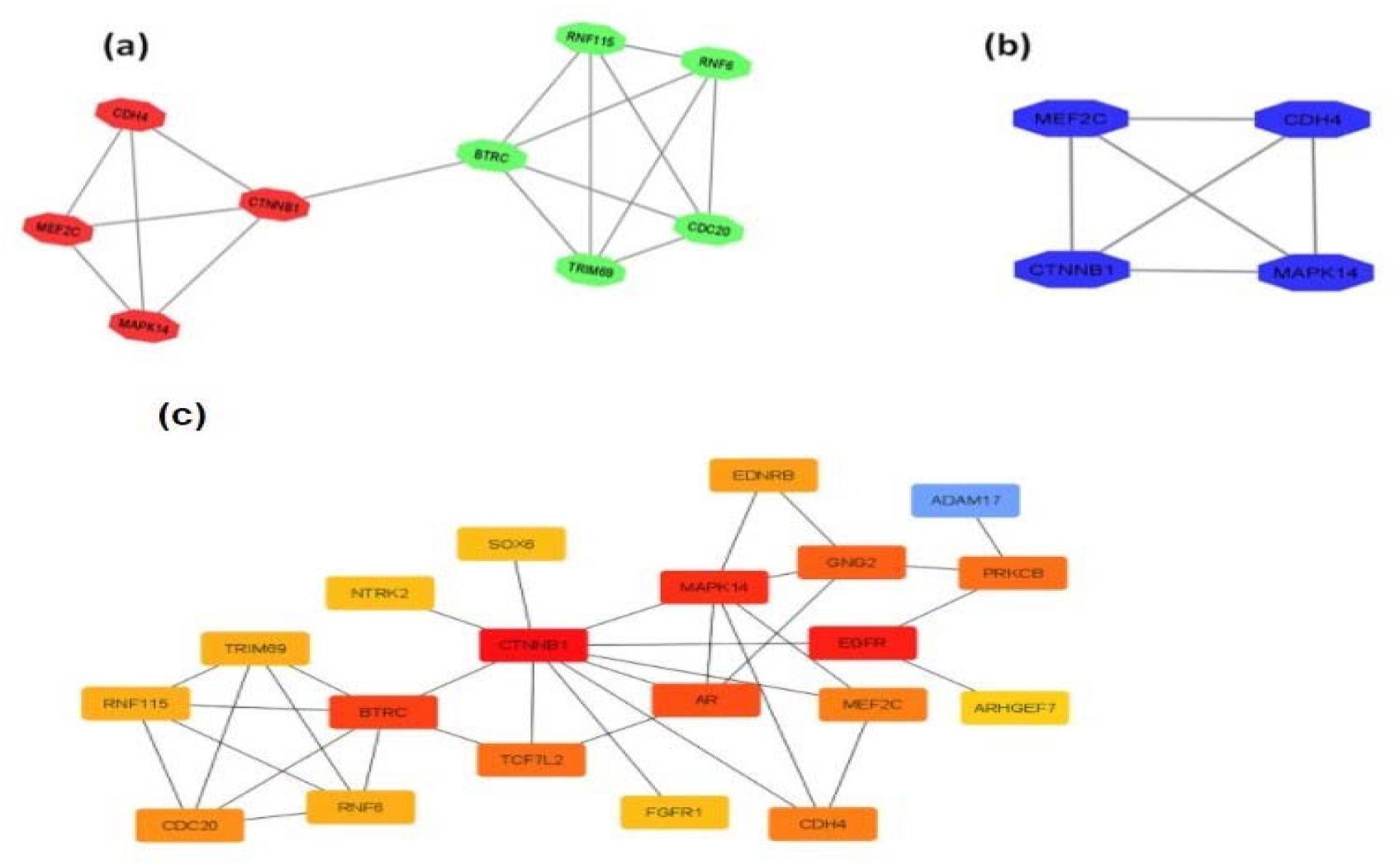
Sub-modules extracted from the PPI network in Cytoscape using the plug-in MCODE; cut off parameters were set as MCODE score >3 and node numbers >4.(a) Sub-module 1 (b) Sub-module 2 (c) Protein - Protein network of the 20 Hub genes using Cytoscape software. The node colors depend on the ranking from 1 to 20 (dark to light).

### Pathway Enrichment analysis of HUB Genes

To investigate the screened HUB genes; pathway enrichment was done using DAVID, KEGG and GO term whereas Heatmaps were structured following which expression of genes was evident. Genes upregulated in both datasets were CTNNB1, CDC20, RNF115; enriched in pathways like-cell differentiation, positive regulation of cell proliferation, mitotic cell cycle, negative regulation of cell death, positive regulation of MAPK cascade, regulation, canonical Wnt receptor signaling pathway, regulation of smooth muscle cell proliferation, Hippo signaling pathway as well as angiogenesis (Table 2). Moreover, genes found to be downregulated in both data sets were-BTRC, CDH4, SOX6, ARHGEF7 which were enriched in the following pathways-regulation of cell size, regulation of growth, cell adhesion, regulation of programmed cell death and positive regulation of gene expression (Table 3). Besides these; EGFR, AR, GNG2, TCF7L2, PRKCB, MEF2C, EDNRB, TRIM69, FGFR1, NTRK2, ADAM17 showed contrasting regulation in both the datasets (Table 2 and 3) and there were also genes like BTRC, MAPK14, EDNRB, RNF6, TRIM69 and FGFR1 that showed no significant change in their levels within the Non-Ocular dataset. (Fig. 4a and 4b) (Table 4)

**Table 2:-.** Upregulated Genes

|  |  |
| --- | --- |
| Genes upregulated in Non-Ocular BCC | EGFR, ADAM17 |
| Genes upregulated in Ocular BCC | AR, GNG2, TCF7L2, PRKCB, MEF2C, EDNRB, TRIM69, FGFR1, NTRK2 |
| Genes upregulated in both | CTNNB1, CDC20, RNF115 |

**Table 3:-.** Downregulated Genes

|  |  |
| --- | --- |
| Genes downregulated in Non-Ocular BCC | AR, GNG2, TCF7L2, PRKCB, MEF2C, EDNRB, TRIM69, FGFR1, NTRK2 |
| Genes downregulated in Ocular BCC | EGFR, MAPK14, RNF6, ADAM17 |
| Genes downregulated in both | BTRC, CDH4, SOX6, ARHGEF7 |

**Table 4:-.** Pathway enrichment of significant genes. (Continuation of this table as Table S1 in Supplement Document)

| TERM | p-Value | Genes |
| --- | --- | --- |
| 30154<br>cell differentiation | 1.6004E-9 | TCF7L2 NTRK2 MEF2C RNF6 EGFR CDC20 AR CDH4<br>ADAM17 EDNRB CTNNB1 SOX6 ARHGEF7 FGFR1 |
| 7166<br>cell surface receptor linked signaling<br>pathway | 1.6014E-8 | TCF7L2 NTRK2 AR ADAM17 EDNRB GNG2 PRKCB<br>CTNNB1 MAPK14 BTRC EGFR FGFR1 |
| 7169<br>transmembrane receptor protein tyrosine<br>kinase signaling pathway | 1.0331E-5 | NTRK2 AR ADAM17 EGFR FGFR1 |
| 60548<br>negative regulation of cell death | 1.0918E-5 | TCF7L2 MEF2C ADAM17 CTNNB1 EGFR FGFR1 |
| 8283<br>cell proliferation | 2.0080E-5 | TCF7L2 AR GNG2 CTNNB1 BTRC EGFR |
| 31657<br>regulation of cyclin-dependent protein<br>kinase activity involved by G1/S | 1.8527E-5 | ADAM17 EGFR |
| 45737<br>positive regulation of cyclin-dependent<br>protein kinase activity | 1.2156E-4 | ADAM17 EGFR |
| 43067<br>regulation of programmed cell death | 1.0754E-4 | TCF7L2 MEF2C ADAM17 CTNNB1 ARHGEF7 EGFR<br>FGFR1 |
| 43410<br>positive regulation of MAPK cascade | 1.9999E-4 | AR CTNNB1 FGFR1 |
| 50678<br>regulation of epithelial cell proliferation | 2.1461E-4 | AR CTNNB1 EGFR |
| 8284<br>positive regulation of cell proliferation | 3.4615E-4 | CDC20 ADAM17 CTNNB1 EGFR FGFR1 |
| 60070<br>canonical Wnt receptor signaling pathway | 1.2012E-3 | TCF7L2 CTNNB1 |
| 48660<br>regulation of smooth muscle cell<br>proliferation | 1.6963E-3 | CTNNB1 EGFR |
| 278<br>mitotic cell cycle | 1.6949E-3 | CDC20 ADAM17 BTRC EGFR |
| 6916<br>anti-apoptosis | 2.5393E-3 | TCF7L2 MEF2C ADAM17 |
| Hsa04912<br>GnRH signaling pathway | 0.01625 | PRKCB, MAPK14, EGFR |
| Hsa04010<br>MAPK signaling pathway | 1.434801984299756E-4 | NTRK2, MEF2C, PRKCB, MAPK14, EGFR, FGFR116 |
| hsa04390<br>Hippo signaling pathway | 0.04168 | TCF7L2, CTNNB1, BTRC |
| Hsa04014<br>Ras signaling pathway | 0.01188 | GNG2, PRKCB, EGFR, FGFR1 |
| 7155<br>cell adhesion | 1.6718E-2 | CDH4 ADAM17 CTNNB1 EGFR |
| 1525<br>angiogenesis | 1.8803E-2 | CTNNB1 FGFR1 |
| 2682<br>regulation of immune system process | 2.0267E-2 | ADAM17 PRKCB CTNNB1 |

**Fig 4 (a).**
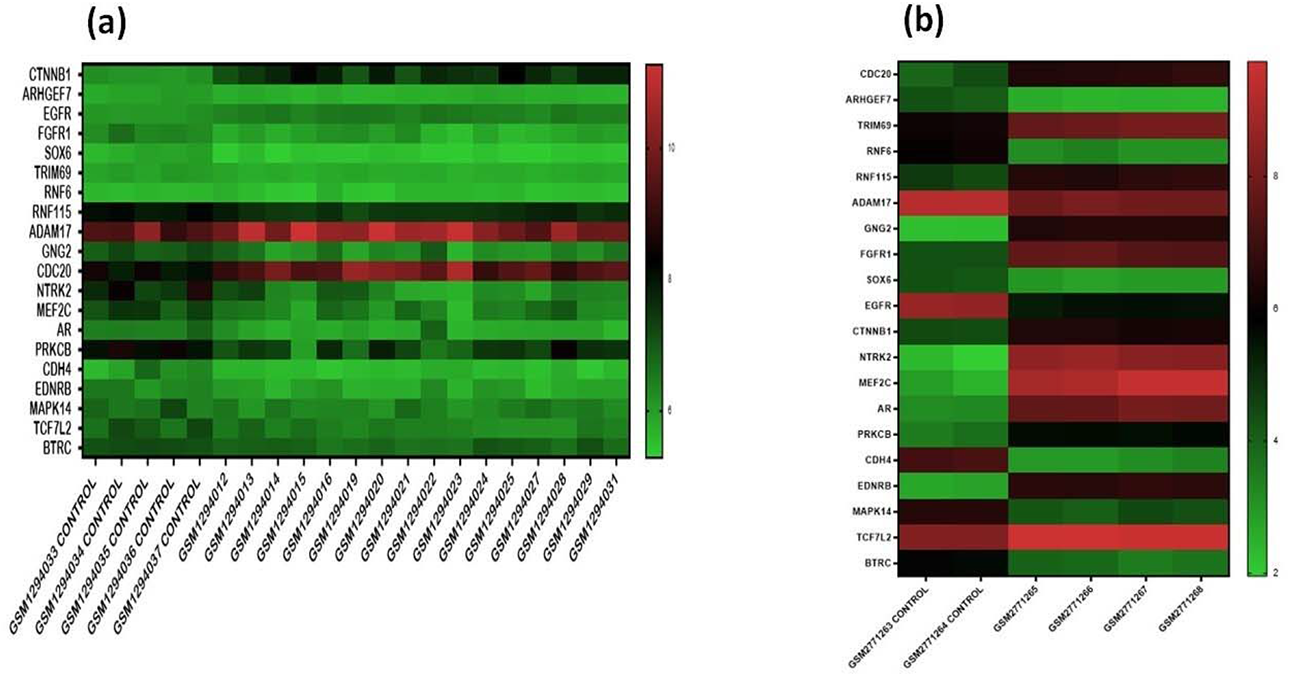
Heatmap showing expression of top 20 significant DEGs in Non-Ocular BCC dataset-GSE53462 compared with suitable control using adj.P ≤ 0.05 threshold. Value of expression progressing from green to red. **(b).** Heatmap showing expression of top 20 significant DEGs in Eyelid BCC dataset-GSE103439 compared with suitable control using adj.P ≤ 0.05 threshold. Value of expression progressing from green to red.

## Discussion

In the last few decades, experimentation on BCC has advanced rapidly specifically concerning the recognition of etiological factors. Many studies have investigated Non-Ocular BCC but rarely focused on BCC of eyelid. Formerly; mRNA profiling of BCC eyelid was accessible but infrequently analysed. In our study, we identified a total of 181 common DEGs out of which 43 DEGs formed a close network in STRING. Reconstructed network in Cytoscape led to the screening of 2 significant sub-network modules via MCODE plugin. 20 HUB genes filtered based on topological parameters like Betweenness Centrality, Degree, Bottleneck and Closeness employing Cytohubba plugin suggested a direction to investigate the complex connection of these genes.

Elucidation of the roles of genes utilizing Pathway enrichment analysis indicated that 59 pathways were closely associated with 20 HUB genes. In particular, it was observed that carcinogenesis and apoptotic pathways were significantly enriched including regulation of programmed cell death, Wnt signaling pathway, Hippo signaling pathway, Ras signaling pathway, cell adhesion and angiogenesis suggesting the role of identified HUB genes in BCC eyelid. The HUB genes with the highest degrees included CTNNB1 which was upregulated in both datasets. CTNNB1 is involved in cell differentiation, negative regulation of cell death, cell proliferation, MAPK cascade, Wnt receptor signaling pathway, Hippo signalling, angiogenesis.^13^ Rajabi et al. demonstrated that it can be used as an IHC marker for the diagnosis of aggressive BCC which is in concurrence with the bioinformatic results obtained.^14^

Another cell cycle regulatory gene, BTRC which is a member of the F-box family, was revealed as a shared DEG downregulated in the both datasets. It plays a key role in mitotic cell cycle, regulation of protein ubiquitination as well as Hippo signaling pathway. Wolter et al. in his studies implicated the role of BTRC in regulation of Wnt pathway in case of cutaneous BCC.^15^ Their findings argued against the major role of BTRC in pathogenesis of cutaneous BCC. Their findings along with our analysis suggest that this gene is of little consequence.

Proto-oncogene, EGFR which belongs to the epidermal growth family is often associated with various cancers. Biray avci et al. in their demonstration has associated the overexpression of EGFR with both clinic and pathogenesis of BCC. This finding does coincide with our analysis in case of Non-Ocular BCC as it was upregulated but its downregulation in BCC eyelid directs us to further investigate the role of EGFR. EGFR does play a role in Hippo signalling. EGFR is reported to be involved in regulation of cyclin-dependent protein kinase activity involved by G1/S, morphogenesis of epithelial fold, regulation of programmed cell death, MAPK signalling and Ras signalling.^16^ Yet another protein-encoding gene, MAPK14 which is a member of the MAP kinase family was upregulated in Non-Ocular BCC but downregulated in BCC eyelid. MAPK14 was associated with Epithelial-Mesenchymal Transition markers in melanomas.^17^

Other genes were found to be concurrently dysregulated in both datasets. One of them being CDC20 which was upregulated in both BCC and is known to interact as well as play a role in the regulation of mitotic cell cycle, in turn controlling the succession of cell proliferation.^18^ The other gene was ARHGEF7 which was mutually downregulated in BCC eyelid and Non-Ocular BCC is reported to play decisive role in the positive regulation of cellular processes, stimulation of Rac1 signalling to focal adhesion as well as seem to function as positive regulator of apoptosis.^19^ Upregulation of CDC20 and downregulation of ARHGEF7 may be involved in the progression and onset of both BCC as suggested by above evidence. GNG2 contributing to cell proliferation, Ras signaling and cell surface receptor signalling showed contrasting differential expression in both the datasets, precisely upregulated in BCC eyelid and downregulated in Non-ocular BCC. It has previously been found to play a role as tumour suppressor in malignant melanomas.^20^ In parallel, Expression of AR differs strikingly in both the datasets with downregulation in Non-Ocular and upregulated in BCC eyelid, its aberrant expression has previously been found in prostate cancer.^21^ Pathways like cell differentiation, regulation of cell size and MAPK cascade regulation shows dysregulation with abnormal AR expressions.

PRKCB which play crucial roles in the regulation of the immune system process, cell surface receptor linked signaling, regulation of gene-specific transcription, cell death, MAPK signaling pathway and Ras signaling pathway showed different expressions in both datasets.^22^ It has been found to be downregulated in renal cell carcinomas whereas upregulated in breast cancers, endometrial cancers and bladder.^23^ Also, ADAM17 showed contrasting expression in both datasets as it was upregulated in Non-Ocular BCC and downregulated in BCC eyelid. This gene is reported to involve in regulation of cell death, cell differentiation as well as cell proliferation in various cancers and enriched in pathways regulating cyclin-dependent protein kinase activity involved by G1/S.^24,25^ Due to their crucial function in these processes, the exact role of the above genes with differing expressions in BCC eyelid and Non-Ocular BCC can be studied further.

Conclusively, our study depicts that text mining and bioinformatic analysis techniques are a useful way to screen genes and understand various pathways responsible for BCC. The molecular events that distinguish these complex biological processes and mRNAs that control the expression of genes at the transcriptional level can be used to develop biomarkers with improved accuracy which is essential for the effective prognosis of BCC eyelid. The study of pathways and genes involved in cell differentiation and various molecular pathways can help us in the early detection and treatment of this melanoma. Common HUB genes involved in the progression of Non-Ocular BCC and BCC eyelid may further be explored and this may help in planning prognostic and therapeutic strategies to cure BCC eyelid employing similar principles that are implemented in Non-Ocular BCC.

## Acknowledgment

We would like to acknowledge Principal, Sri Venkateswara College, University of Delhi for her support.

## Supporting Information

FIGURE S1 and TABLE S1 (continuation of Table 4) is submitted as supplement file.

## Notes

https://www.ncbi.nlm.nih.gov/geo/

### Competing Interest Statement

The authors have declared no competing interest.

### Summary of Updates

Author order updated

