## Supplementary material for "Identification of Differentially Expressed Genes and associated pathways common to Eyelid and Non-Ocular Basal Cell Carcinoma to understand the Molecular Biology of BCC": Fig S1 and Table S1

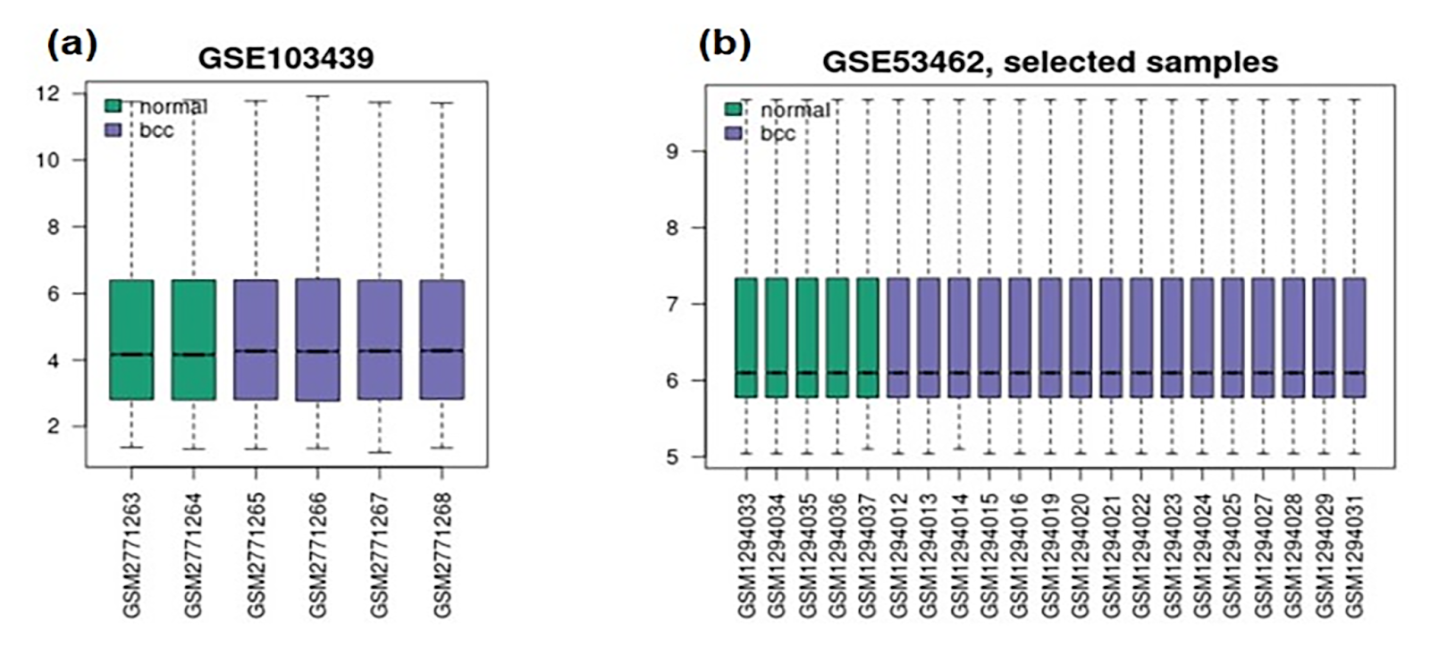

**Fig S1**. The quality of expression data. The datasets were normalised and the expression depicted that the datasets (a) GSE103439 and (b) GSE53462 were of high quality. a) The normalised data of Eyelid BCC b) The normalised data of Non-Ocular BCC.

| TERM | p-Value | Genes |
| --- | --- | --- |
| 48869  cellular developmental process | 2.2988E-9 | TCF7L2 NTRK2 MEF2C RNF6 EGFR CDC20 AR CDH4 ADAM17 EDNRB CTNNB1 SOX6 ARHGEF7 FGFR1 |
| 48522  positive regulation of cellular process | 1.8050E-8 | TCF7L2 MEF2C PRKCB RNF6 EGFR CDC20 AR CDH4 ADAM17 CTNNB1 BTRC SOX6 ARHGEF7 FGFR1 |
| 31325  positive regulation of cellular metabolic process | 1.8486E-7 | CDC20 TCF7L2 AR MEF2C ADAM17 CTNNB1 RNF6 BTRC SOX6 EGFR |
| 32583  regulation of gene-specific transcription | 3.2185E-7 | TCF7L2 AR MEF2C PRKCB CTNNB1 RNF6 |
| 10909  positive regulation of heparan sulfate proteoglycan biosynthetic process | 1.8574E-6 | TCF7L2 CTNNB1 |
| 90066  regulation of anatomical structure size | 3.3698E-6 | AR CDH4 ADAM17 EDNRB RNF6 FGFR1 |
| 42063  gliogenesis | 2.2663E-6 | CTNNB1 SOX6 ARHGEF7 EGFR |
| 10604  positive regulation of macromolecule metabolic process | 1.9097E-6 | CDC20 TCF7L2 AR MEF2C ADAM17 CTNNB1 RNF6 BTRC SOX6 |
| 8361  regulation of cell size | 1.0331E-5 | AR CDH4 ADAM17 RNF6 FGFR1 |
| 7169  transmembrane receptor protein tyrosine kinase signaling pathway | 1.0331E-5 | NTRK2 AR ADAM17 EGFR FGFR1 |
| 1763  morphogenesis of a branching structure | 2.1256E-5 | AR CTNNB1 BTRC FGFR1 |
| 31659  positive regulation of cyclin-dependent protein kinase activity involved in G1/S | 1.8527E-5 | ADAM17 EGFR |
| 9653  anatomical structure morphogenesis | 1.6162E-5 | TCF7L2 AR CDH4 EDNRB CTNNB1 BTRC SOX6 EGFR FGFR1 |
| 44093  positive regulation of molecular function | 1.5838E-5 | CDC20 TCF7L2 NTRK2 ADAM17 EDNRB BTRC EGFR |
| 48468  cell development | 1.4893E-5 | TCF7L2 CDH4 EDNRB CTNNB1 SOX6 EGFR FGFR1 |
| 45165  cell fate commitment | 3.3657E-5 | TCF7L2 CTNNB1 SOX6 FGFR1 |
| 60571  morphogenesis of an epithelial fold | 1.4354E-4 | AR EGFR |
| 10628  positive regulation of gene expression | 1.2783E-4 | TCF7L2 AR MEF2C CTNNB1 RNF6 SOX6 |
| 48519  negative regulation of biological process | 1.4680E-4 | CDC20 TCF7L2 AR MEF2C ADAM17 CTNNB1 RNF6 SOX6 EGFR FGFR1 |
| 45737  positive regulation of cyclin-dependent protein kinase activity | 1.2156E-4 | ADAM17 EGFR |
| 40008  regulation of growth | 1.1003E-4 | AR CDH4 ADAM17 RNF6 FGFR1 |
| 65008  regulation of biological quality | 1.0657E-4 | TCF7L2 AR CDH4 ADAM17 EDNRB CTNNB1 RNF6 SOX6 FGFR1 |
| 50678  regulation of epithelial cell proliferation | 2.1461E-4 | AR CTNNB1 EGFR |
| 45445  myoblast differentiation | 2.8039E-4 | TCF7L2 CTNNB1 |
| 31323  regulation of cellular metabolic process | 2.9156E-4 | TCF7L2 NTRK2 MEF2C PRKCB RNF6 EGFR CDC20 AR ADAM17 EDNRB CTNNB1 BTRC SOX6 |
| 50789  regulation of biological process | 3.5805E-4 | TCF7L2 NTRK2 MEF2C PRKCB RNF6 MAPK14 EGFR CDC20 AR CDH4 ADAM17 EDNRB CTNNB1 BTRC SOX6 ARHGEF7 FGFR1 |
| 50789  regulation of biological process | 3.5805E-4 | TCF7L2 NTRK2 MEF2C PRKCB RNF6 MAPK14 EGFR CDC20 AR CDH4 ADAM17 EDNRB CTNNB1 BTRC SOX6 ARHGEF7 FGFR1 |
| 2053  positive regulation of mesenchymal cell proliferation | 3.8388E-4 | CTNNB1 FGFR1 |
| 32989  cellular component morphogenesis | 1.2782E-3 | CDH4 CTNNB1 SOX6 EGFR |
| 48660  regulation of smooth muscle cell proliferation | 1.6963E-3 | CTNNB1 EGFR |
| 278  mitotic cell cycle | 1.6949E-3 | CDC20 ADAM17 BTRC EGFR |
| 43170  macromolecule metabolic process | 1.0169E-2 | CDC20 NTRK2 AR ADAM17 RNF115 PRKCB RNF6 MAPK14 BTRC EGFR FGFR1 |
| 30817  regulation of cAMP biosynthetic process | 1.0711E-2 | NTRK2 EDNRB |
| 31396  regulation of protein ubiquitination | 1.2207E-2 | CDC20 BTRC |
| 19219  regulation of nucleobase, nucleoside, nucleotide and nucleic acid metabolic process | 1.3984E-2 | TCF7L2 NTRK2 AR MEF2C EDNRB PRKCB CTNNB1 RNF6 SOX6 |
| Hsa04912  GnRH signaling pathway | 0.01625 | PRKCB, MAPK14, EGFR |
| hsa04390  Hippo signalling | 0.04168 | TCF7L2, CTNNB1, BTRC. |
| 8283  cell proliferation | 2.0080E-5 | TCF7L2 AR GNG2 CTNNB1 BTRC EGFR |
| 32879  regulation of localization | 1.6718E-2 | TCF7L2 ADAM17 CTNNB1 EGFR |
| 30318  melanocyte differentiation | 1.8033E-2 | EDNRB |
| 70201  regulation of establishment of protein localization | 1.9036E-2 | TCF7L2 EGFR |
| 2682  regulation of immune system process | 2.0267E-2 | ADAM17 PRKCB CTNNB1 |

**TABLE S1 : Pathway enrichment of significant genes (Continuation of Table 4)**
